# Structural visualization of de novo initiation of RNA polymerase II transcription

**DOI:** 10.1101/2021.05.03.442346

**Authors:** Chun Yang, Rina Fujiwara, Hee Jong Kim, Jose J. Gorbea Colón, Stefan Steimle, Benjamin A. Garcia, Kenji Murakami

## Abstract

Structural studies of the initiation-elongation transition of RNA polymerase II (pol II) transcription were previously facilitated by the use of synthetic oligonucleotides. Here we report structures of initiation complexes *de novo* converted from pre-initiation complex (PIC) through catalytic activities and stalled at different template positions. Contrary to previous models, the closed-to-open promoter transition was accompanied by a large positional change of the general transcription factor TFIIH which became in closer proximity to TFIIE for the active delivery of the downstream DNA to the pol II active center. The initially-transcribing complex (ITC) reeled over 80 base pairs of the downstream DNA by scrunching, while retaining the fixed upstream contact, and underwent the transition to elongation when it encountered promoter-proximal pol II from a preceding round of transcription. TFIIH is therefore conducive to promoter melting, TSS scanning, and promoter escape, extending far beyond synthesis of a short transcript.

## Introduction

RNA polymerase II (pol II) and the six general transcription factors (GTFs) assemble in a transcription pre-initiation complex (PIC), which recognizes promoter DNA before every round of transcription, and opens the double-stranded DNA to expose and select a transcription start site (TSS) (Conaway and Conaway, 1993; Kornberg, 2007). Following the TSS recognition, the PIC transitions to the initially-transcribing complex (ITC), which is responsible for synthesizing a nascent transcript and subsequently transitions to an elongation complex (EC), followed by re-initiation. This set of transitions is universal across all eukaryotes and overlaid by many additional regulatory steps involving elongation factors such as DSIF (Spt4/5 in yeast) and initiation factors such as Mediator (Adelman and Lis, 2012; Conaway and Conaway, 2012; Wade and Struhl, 2008). It is also commonly thought that polymerases from successive rounds of transcription are located immediately adjacent to each other at promoter proximal regions of actively transcribed genes, yet the precise nature and the significance of the interaction remain unknown (Ehrensberger et al., 2013).

The largest GTF, TFIIH comprising 10 subunits, is an integral component of the PIC: the translocase subunit Ssl2 (XPB in humans) acts as a molecular motor during promoter opening, TSS scanning, and initial RNA chain elongation (Bradsher et al., 2000; Dvir et al., 1997; Fazal et al., 2015; Fishburn et al., 2015; Qiu et al., 2020; Spangler et al., 2001). The other subunits comprise the six-subunit structural core (Greber et al., 2019) and the three-subunit kinase termed TFIIK for pol II CTD phosphorylation (van Eeuwen et al., 2021a). Previous structural studies of open promoter complexes provided information about locations of GTFs and the DNA path (He et al., 2013; He et al., 2016; Schilbach et al., 2017). However the open promoter template was not obtained by the catalytic activity of TFIIH. Thus it remains to be determined how TFIIH is responsible for the initiation process.

To elucidate the mechanisms of the initiation, we have recently developed an in vitro transcription system, in which pol II and six GTFs (TFIIA, TBP, TFIIB, TFIIE, TFIIF, and TFIIH) isolated from the yeast *Saccharomyces cerevisiae*, melt double-stranded promoter DNA, and initiate RNA synthesis de novo with high efficiency (Fujiwara and Murakami, 2019; Murakami et al., 2013a; Murakami et al., 2015a). Resulting post-initiation complexes could be stalled at different template positions on a series of G-less promoter mutants and isolated in abundant homogeneous form by glycerol gradient sedimentation (Fujiwara et al., 2019).

Here we report cryo-EM structures of such de novo initiation complexes stalled at two different template positions (Figure 1A). The first cryo-EM structure with the G-less 26 template revealed an ITC containing all the GTFs, pol II, and a nascent RNA on a bona fide open promoter DNA. Compared to previous structures of PICs (Dienemann et al., 2019; Murakami et al., 2015b; Schilbach et al., 2017), the ITC underwent a large positional change in TFIIH for the active delivery of the downstream DNA to the pol II active center. By contrast, the second structure with the G-less 49 template revealed successive elongation complexes (EC+EC), in which two polymerases that completed promoter escape were in close contact with each other. From the combination of these structures with previous biochemical and biophysical studies (Fazal et al., 2015; Fujiwara et al., 2019), we arrive at a picture of the initiation process driven by TFIIH, in which the preceding pol II stalled at promoter-proximal regions blocks TFIIH translocation of the trailing ITC and ultimately occludes TFIIH on a promoter.

**Figure 1.**
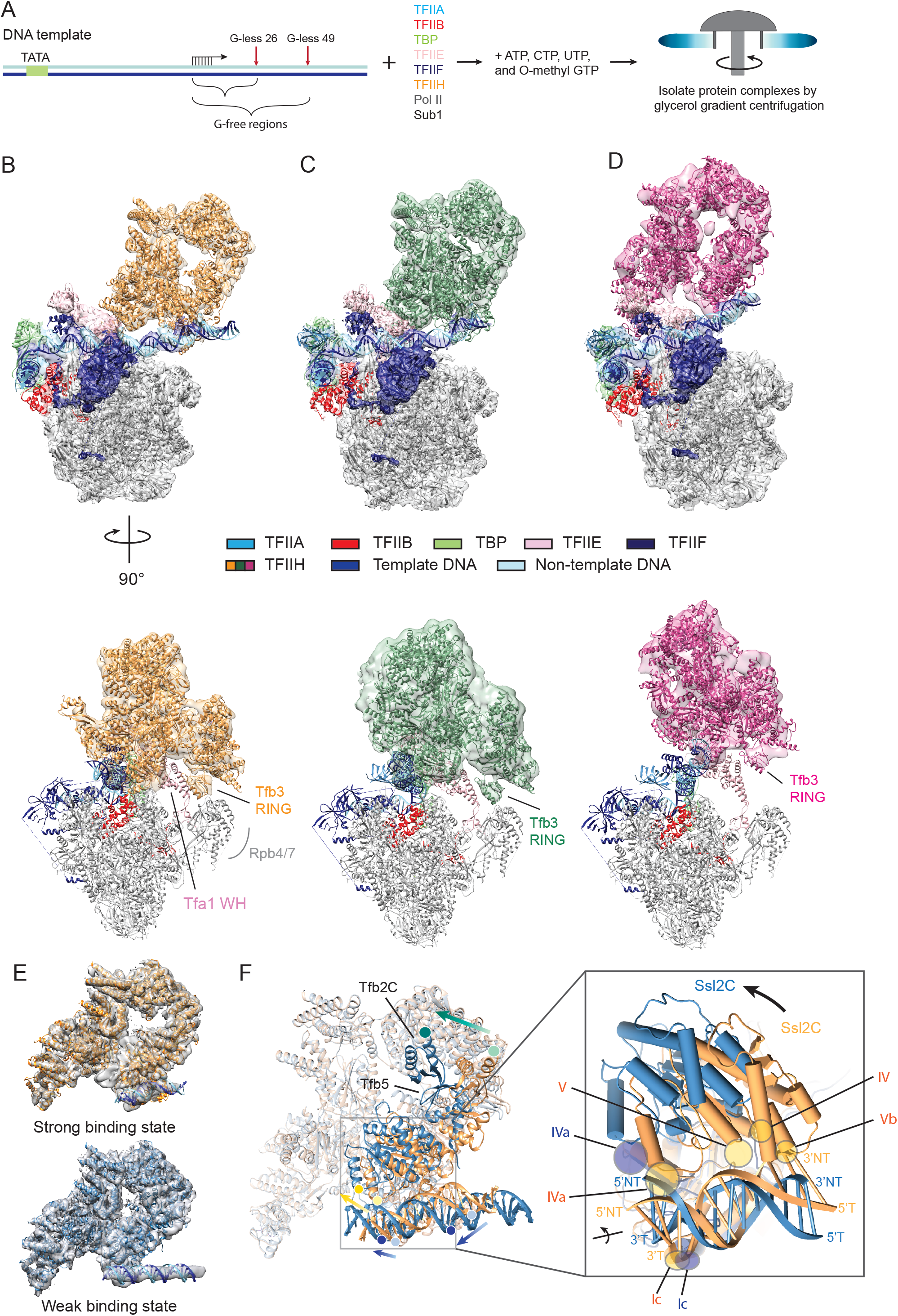
Structures of three forms of pre-initiation complexes. **(A)** Schematic representation for isolation of the transcription complexes analyzed in this study. **(B-D)** Composite density maps and the models of three forms of PIC on G-less 26 DNA template; PIC1 (B), PIC2 (C), and PIC3 (D). Side view (top) and front view (bottom) are shown. Same colors are used throughout the manuscript unless otherwise noted: TFIIA (steel blue), TFIIB (red), TBP (light green), TFIIE (pink), TFIIF (dark blue), TFIIH (orange, green, or magenta), template DNA (blue), non-template DNA (sky blue). Tfb3 interacts with Rpb4/7 in PIC1 and PIC2, but dissociates in PIC3. **(E)** Composite density maps and models of strong (top) and weak (bottom) binding states of TFIIH, with corresponding models in orange and steel blue, respectively. **(F)** Comparison of TFIIH and downstream DNA in the strong and weak binding states. Same residues of Tfb2C, Ssl2C, and DNA in the two states are marked by circular dots and the directions of movements are indicated by green, yellow, and blue arrows, respectively. Inset, Ssl2–DNA interactions suggested by the model. The five DNA binding motifs (Ic, IVa, IV, V, Vb) are indicated.

## Results

### Isolation of bona fide ITCs and Cryo-EM analysis

ITCs were obtained by transcription reaction with the G-less 26 SNR20 promoter fragment, in the procedure we have previously established (Fujiwara et al., 2019) (Figure 1A). Briefly the protocol entails combining a 33-subunit PIC with a G-less SNR20 promoter fragment, supplemented with ~4-8 fold molar excess pol II and GTFs relative to PIC, followed by addition of NTPs for transcription reaction. ITCs were stalled at position +26 relative to the TSS (+1) by use of chain-terminating 3’-O-methyl GTP instead of normal GTP, while inclusion of 4’-thio UTP instead of normal UTP induced pol II arrest and thereby prevented the extensive backtracking, which would otherwise have completely collapsed back to the closed complex (PIC) with concomitant RNA release during subsequent gradient sedimentation (Fujiwara et al., 2019). The reaction mixture was sedimented on a 10-40% of glycerol gradient to remove free nucleotides and excess GTFs and pol II. The resulting ITC contained equimolar amounts of GTFs and pol II, and transcripts of ~20-26 nucleotides initiating from positions +1 to +7 (Figures S1A-B). Due to the similarity in size, ITC were not separable on the gradient from residual PICs that did not engage in transcription and/or those that collapsed back from ITC.

Knowing the heterogeneity of the specimen, aliquots of peak fractions were embedded in ice, disclosing fields of monodispersed particles (Figure S1C). We imaged ~4 million particles using Titan Krios electron microscopes equipped with a K3 direct electron detector. 2D class averaging of particles yielded a set of homogeneous classes, which showed clear division in two parts: a well-ordered pol II and a disordered TFIIH (Figure S1D). For some classes, DNA was identifiable on TFIIH. We selected a subset of particles (~0.8 million) through 2D class averaging and subjected them to *ab initio* calculation of an initial map (Figure S1E). To sort out variability in positions of TFIIH and DNA, the ~1.8 million particles were subjected to iterative global 3D classifications, which revealed three forms of PICs (hereinafter PIC1, PIC2, PIC3) and one form of the ITC, accounting for 137K, 117K, 69K, and 120K particles, respectively. In each form of PICs and ITC, TFIIH and DNA were poorly ordered due to their flexibility compared to pol II. For reconstruction of PIC1-3, TFIIH was subjected to focused classification and refinement, and composited back to the entire map (Figure S1F-N). For reconstruction of the ITC, three masks were created, the first containing the active center of pol II, the second containing TFIIH, and the third containing DNA-TFIIA-TBP-TFIIE (Tfa1 and Tfa2 WH domains)-TFIIF (Tfg2 WH domains)-TFIIB (cyclin domains). Three segments were subtracted from images with respective masks, subjected to local 3D classifications and refinement, and then composited back to the entire complex (Figure S1O-Q). Focused classification of pol II active center in ITC map enabled removal of particles that had only poor density of the DNA-RNA hybrid.

Three forms of PICs and the ITC differed from each other in locations and conformations of TFIIH and DNA path (Figures 1B-D). PIC1 was a good match to previous structures of yeast 31-subunit PIC (EMDB 3114 and EMDB0092) (Dienemann et al., 2019; Murakami et al., 2015b), and was resolved at higher resolution (3.2-4.6 Å) than before, attesting to our sample preparation and data analysis strategy. In PIC1, TFIIH was resolved at near atomic resolution (4.6 Å), allowing us to define two different DNA-binding modes for DNA translocation, as described in detail below. PIC2 and PIC3 were refined to 3.2-7.3 Å and 3.4-11.8 Å, respectively, and were distinct from PIC1 in the position and the conformation of TFIIH and DNA. The ITC was refined to 3.2-9.9 Å resolution, revealing a bona fide open promoter DNA and a short 6-bp DNA-RNA hybrid in the pol II active center, which differs from previous open promoter complexes with artificial templates (He et al., 2016), by translations of more than ~30 Å (over 50 Å for some TFIIH subunits (Tfb1, Tfb2, Tfb4, Tfb5)) in the location of TFIIH.

### Two DNA-binding modes of TFIIH in PIC1

The previous cryo-EM model of PIC (EMDB 3114 and EMDB0092) was well fitted into density of PIC1. The fit showed some differences in degree and position of the DNA bend ~20-30 bp downstream of the TATA box, which may relate to different promoter sequences (SNR20 in this study vs HIS4 in previous studies). The relationship between the DNA bend/distortion and promoter melting was previously characterized (Dienemann et al., 2019). The model of TFIIH was built using the previous 3.9 Å-resolution cryo-EM structure of yeast TFIIH bound to Rad3-Rad23-Rad33 as an initial model (van Eeuwen et al., 2021b). The model of PIC1 was subjected to iterative refinements with Coot (Emsley et al., 2010) and Phenix (Liebschner et al., 2019) (Supplemental Table 1), and then used as an initial template for model building of PIC2, PIC3 and ITC.

Focused 3D classification of TFIIH in PIC1 revealed two forms of TFIIH at 4.6 Å and 7.6 Å resolution (orange vs steel blue in Figures 1E-F, Figure S1F). One form had a good match to the previous structure of the pre-translocation state of TFIIH in the PIC (Dienemann et al., 2019; Murakami et al., 2015b; Schilbach et al., 2017) (orange, upper panel of Figure 1E), while the other form revealed a ~60° rotation of the domain that consists of Tfb5 and the C-terminal region of Tfb2, accompanied by a rotation of the C-terminal ATPase domain of Ssl2 (Ssl2C) relative to the rest of TFIIH (steel blue, lower panel of Figure 1E) as previously observed in the structure of TFIIH-Rad4-Rad23-Rad33 (van Eeuwen et al., 2021b). In the former (orange in Figures 1E-F), a ~13-bp segment of DNA double helix was bent, deep within the DNA-binding groove between the two ATPase domains, in close contact with the five DNA binding motifs (Ic, IVa, IV, V, Vb, as previously defined (Fairman-Williams et al., 2010)) (referred to as strong-binding state), whereas, in the latter (steel blue in Figures 1E-F), the DNA was relatively straight, only in contact with two DNA binding motifs (IVa, Ic) (referred to as weak-binding state). In the weak-binding state, the detachment of the DNA from motifs IV, V, and Vb was accompanied by the rotation of Ssl2C along with Tfb5-Tfb2C (Figure 1F), consistent with the previously suggested role of Tfb5 (p8 in humans) in stimulating Ssl2’s catalytic activity (Coin et al., 2006; Ranish et al., 2004). The remaining DNA-Ssl2 interactions by the two motifs IVa and Ic were altered, enabling a slight rotation of the DNA along its axis, likely coupled with DNA translocation (Figure S1R). Thus the weak-binding state may represent the post-translocation state, although nucleotides were not directly resolved.

### Distinct forms of PICs represent the path to the open promoter complex

PIC2 and PIC3 differed from PIC1 in locations and conformations of TFIIH and DNA path, as readily apparent in initial rounds of 3D classification (Figure S1E), and there are several notable differences between three forms of the PIC (Figure 2). First, PIC2 and PIC3 differed from PIC1 by ~20 Å and ~30 Å shifts in the location of TFIIH (Figure 2A), and by repositioning of Ssl2 on DNA by one turn of dsDNA (~10 bp), accompanied by greater degrees of DNA distortion ~20-30 bp downstream of the TATA box (Figures 2B-C). Second, PIC2 and PIC3 revealed TFIIH in the weak-binding state, while PIC1 primarily revealed the strong-binding state, suggesting that locations of TFIIH in PICs would shift the conformational equilibrium among coexisting translocation states. Third, in PIC3, Tfa1 (TFIIE) and the RING domain of Tfb3 (one of three TFIIK subunits) were dissociated from the pol II clamp and Rpb4/7, resulting in a shift in their positions by ~10 Å relative to those in PIC1/PIC2, such that TFIIH and TFIIE less closely contacted pol II (lower panels of Figures 1B-D). Our previous exonuclease footprinting demonstrated that omission of TFIIK, while retaining a high-level of TFIIK-independent transcription, causes upstream shift of the downstream boundary of the PIC (by ~5 residues) (Murakami et al., 2015a), suggesting removal of TFIIK may be able to mimic the transition to PIC3. Irrespective of these significant differences between three forms, promoter DNA was nevertheless associated only with GTFs and not with pol II in all forms, requiring for the translocase activity of TFIIH for promoter melting.

**Figure 2.**
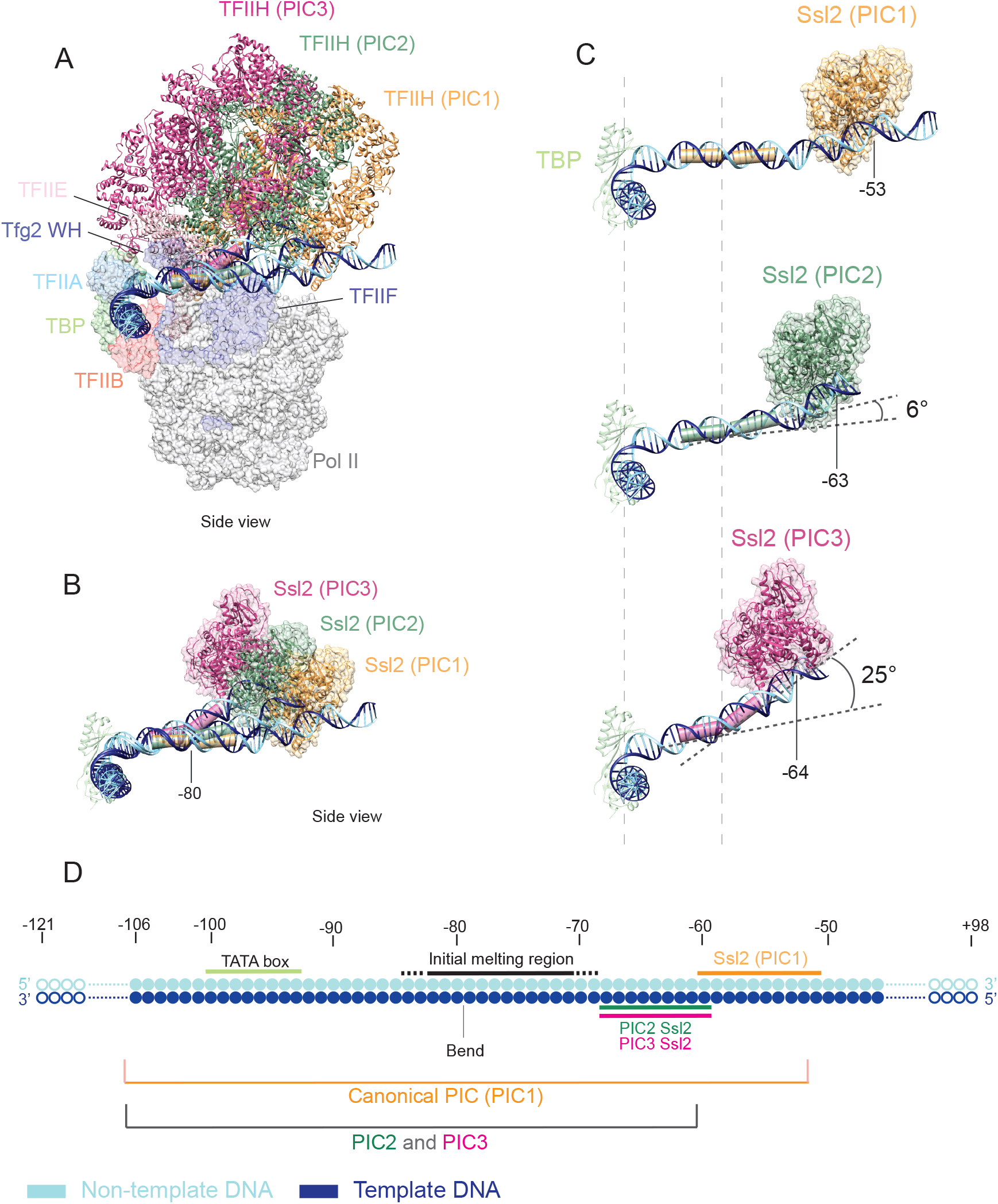
Distortion of promoter DNA in PIC1-3. Coloring as in Figure 1. **(A)** Comparison of locations of TFIIH and DNA in PIC1-3 relative to Pol II. TFIIH shifts ~ 20 Å and ~30 Å in PIC2 and PIC3, respectively, relative to that in PIC1. **(B)** Paths of promoter DNA. Ssl2 contacting DNA is shown. **(C)** Ssl2 binds ~47 bp and ~37 bp downstream of the TATA box in PIC1 and PIC2/PIC3, respectively. DNA bends by ~6° and ~25° in PIC2 and PIC3, respectively. Dashed lines indicate positions of the bend at ~–80 and the TATA box at –100. The numbering is relative to the TSS. **(D)** A schematic showing PIC1-3 occupancy on the promoter DNA.

### Bona fide ITC structure

Locations of GTFs in the ITC largely correspond to those in the PIC3 except some differences in orientations of TFIIH and TFIIE (Figure 3A, Movie S1); as in PIC3, the Tfb3 RING domain was absent on Rpb4/7 (not visualized in the map), so that TFIIH and TFIIE less closely contacted pol II (Figure 3A). The promoter DNA of the ITC was suspended above the pol II cleft, bound by TFIIH at the downstream end, and by the remaining general transcription factors (GTFs) at upstream end. A ~36 bp segment (positions –116 to –79) of the upstream DNA bound to TFIIA, TBP, TFIIB (cyclin domains), TFIIE (Tfa1 and Tfa2 WH1 and WH2 domains), and TFIIF (Tfg2 WH domains) was clearly discerned (Figure 3B). The upstream edge of the transcription bubble in the ITC (position –79), corresponding to the position of the 25° bend in the PIC3, was stabilized by the WH domain of Tfa1 (the large subunit of TFIIE) (Figure 3B). Although the downstream DNA bound to Ssl2 was poorly ordered, there was a discernable density attributable to a ~9-bp short segment of DNA double helix bound to Ssl2 in the focused classification of TFIIH (Figures S3A-B). In between, the DNA of over ~100 bp was missing except the region of the DNA (–2 to +9) that was accommodated in the pol II active center (Figure 3C): the region of the DNA between positions –79 and –2 presumably looped out or scrunched (Fazal et al., 2015; Kapanidis et al., 2006; Liu et al., 2010), while the downstream DNA that bridges between Ssl2 and the pol II active site, likely a straight DNA double helix, was disordered (schematically illustrated in Figure 3E). It is important to note that the downstream DNA was not observed deep in pol II downstream cleft, which markedly contrasts to the transcribing complex (EC) (Gnatt et al., 2001; Kettenberger et al., 2004) (inset of Figure 3E).

**Figure 3.**
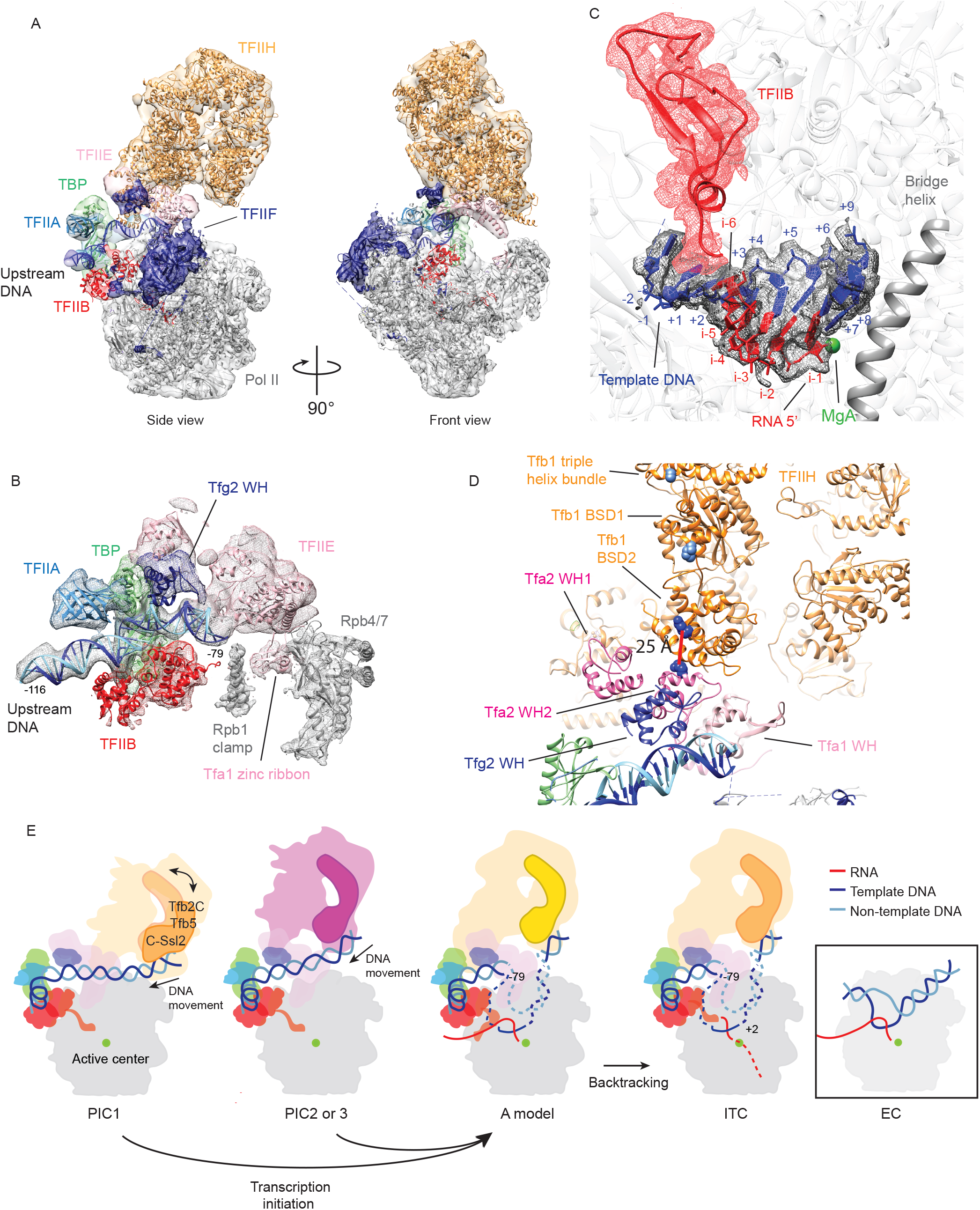
Structure of ITC with the G-less 26 template. **(A)** Cryo-EM map and a corresponding model of the ITC. **(B)** EM density map shows the DNA-RNA hybrid in the pol II active center in contact with TFIIB. **(C)** EM density shows the upstream DNA bound to TFIIA, TBP, TFIIB (cyclin domains), TFIIE (Tfa1 and Tfa2 WH1 and WH2 domains), and TFIIF (Tfg2 WH domains). **(D)** TFIIE-TFIIH interactions in the ITC. Tfb1 BSD2 domain (orange) is in contact with Tfa2 WH2 (hot pink). Red line indicates a cross-link between K268 of Tfb1 and K194 of Tfa2. K581 of the triple helix bundle and K179 of the BSD1 (sky blue) forms cross-links with the C-terminal region of Tfa1. **(E)** Schematic of transition from PICs to ITC. Inset, schematic of EC viewed from the same orientation as PIC1-3 and ITC.

The short DNA-RNA hybrid observed in the active center of the ITC was a good match to the 6-bp DNA-RNA hybrid previously observed by X-ray crystallography in a complex with TFIIB (Sainsbury et al., 2013) (Figure 3C): eleven nucleotides of the template strand at positions –2 to +9 were identifiable with discernible backbone phosphate positions. The six ribonucleotides of RNA in the hybrid were identifiable at positions from i–1 to i–6 relative to the nucleotide addition site, i+1. The 5’-terminal nucleotide (position i–6) was in direct contact with the finger domain of TFIIB (Figure 3C). Despite of inclusion of 4’-thio UTP, pol II was evidently subjected to extensive backtracking from +26 to +6, so that the RNA was stabilized in the ITC, that would otherwise have been incompatible with TFIIB (Bushnell et al., 2004) (Figure 3E). Consistent with this model, there was a density attributable to the backtracked RNA (at positions i+3 to i+5) in the pol II funnel. The observed path of the backtracked RNA coincides with that of the backtracked EC (see below).

As a key feature of the ITC, the positional constraint of TFIIH imposed by the rigid straight double-stranded DNA (as in PIC1) as well as by the contact between the Tfb3 RING domain and Rpb4/7/Tfa1 was relieved due to the promoter melting, such that TFIIH was stabilized through protein-protein interactions that were absent in the canonical PIC (PIC1) (Figure 3D, Figure S3, Movie S1): the primary contact was made between the Tfb1 BSD2 domain of TFIIH and the Tfa1 WH2 domain of TFIIE. The second contact was made between the Rad3 Arch domain of TFIIH and the Tfa2 WH2 domain of TFIIE (Movie S1). Although not modeled, there was a density adjacent to the Tfb1 BSD1 domain of TFIIH, which may be attributed to the C-terminal region of Tfa1. These interactions are in good agreement with a number of cross-links, most of which were obtained in the PIC lacking TFIIK (Murakami et al., 2013b) (e.g., a cross-link between K268 of Tfb1 and K194 of Tfa2, Figure 3D and S3C-F).

The TFIIH-TFIIE interactions described above (Figure 3D) apparently serve as a critical point of contact between TFIIH and the remaining GTFs, such that TFIIH rotates the downstream DNA for unwinding, while retaining fixed upstream contact (Fishburn et al., 2015; Kim et al., 1997). Without TFIIH being held by this anchor point, TFIIH itself may freely rotate around the DNA axis. Based on real-time observations of single PICs (Fazal et al., 2015), this translocation reels dozens of base pairs of the downstream DNA independently of pol II transcription (*i.e.,* only with dATP that allows for DNA translocation by Ssl2), and continues even after the point (~+7–+12) at which TFIIB is displaced from the RNA exit tunnel, in good agreement with biochemical isolation of stable ITCs stalled at ~+26–27 (Fujiwara et al., 2019).

### The structure of ECs colliding head-to-end (EC+EC)

In contrast to the G-less 26 complex that formed such long-persisting ITC, our previous biochemical studies demonstrated that the G-less 49 complex contained a pol II that escaped the promoter (+49), and another pol II that initiated transcription by re-utilizing the promoter to generate the ~25 nt RNAs (thus referred to as re-initiation complex) (Fujiwara et al., 2019). Upon removal of ATP during gradient sedimentation, the ~25 nt transcripts from the second round of transcription were retained in the G-less 49 complex, in contrast to the G-less 26 complex that released transcripts of similar lengths by extensive backtracking of pol II (Fujiwara et al., 2019). Inclusion of 4’-thio-UTP, instead of normal UTPs, was needed to induce pol II arrest and prevent RNA release for the structure determination of the ITC with the G-less 26 template, as described above. This indicates that an EC stalled at promoter proximal regions (~+49) serves to play a positive role in the trailing ITC, as previously suggested (Ehrensberger et al., 2013).

To isolate G-less 49 complexes for structural study, transcription reaction with the G-less 49 template was initiated by adding NTPs (ATP, CTP, and UTP) with chain-terminating 3’-O-methyl GTP, and following gradient sedimentation revealed two major peaks of the re-initiation complex (fractions 17-18 and 22-24 in Figure 4A). Aliquots of each peak were subjected to cryo-EM analysis in a similar manner to the G-less 26 complex (Figure S4). Consistent with protein analysis by SDS-PAGE (Figure 4B), initial two rounds of 2D classification of the slower sedimenting fractions yielded a set of well-ordered homogeneous classes of two colliding pol II molecules (referred to as EC+EC) (Figures 4C and 4E), while the faster sedimenting fractions yielded similar classes of two colliding pol II molecules associated with a set of GTFs (referred to as EC+ITC) (Figures 4D and 4F). After interactive 3D classifications to remove some residual PICs (Figure S4F, see also Figure 4B), the structure of the EC+EC was refined to 3.5 Å resolution (Figure 5A), while the considerable variability in the distance between two pol II molecules prevented refinement of EC+ITC past about 15-Å resolution.

**Figure 4.**
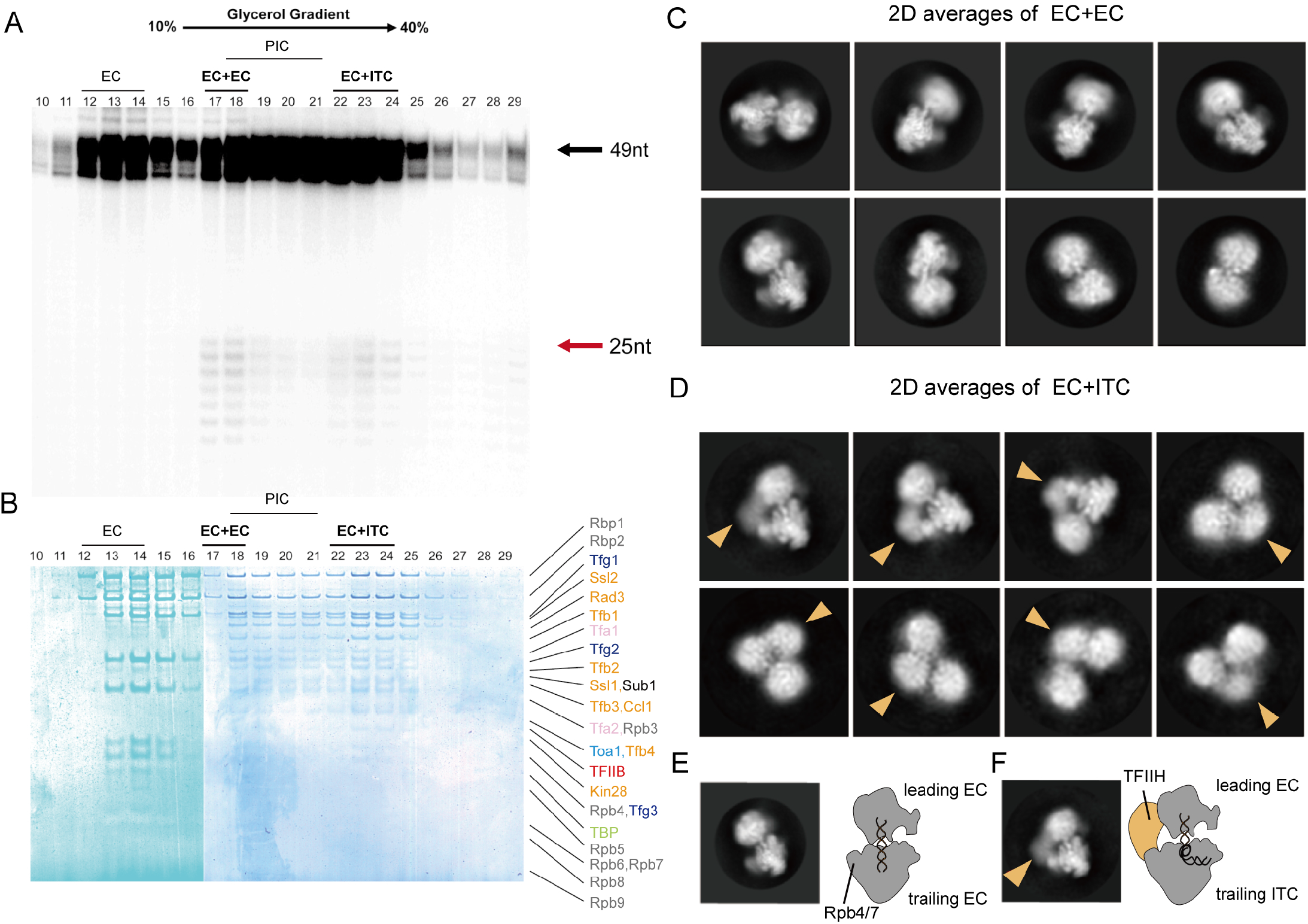
G-less 49 complexes contain two major post-initiation complexes, EC+EC and EC+ITC. **(A-B)** Transcription complexes with the G-less 49 template were subjected to glycerol gradient sedimentation. RNA analysis of the fractions by denaturing Urea-PAGE gel (A) and protein analysis of the fractions by SDS-PAGE gel (B) revealed EC+EC (fractions 17-18) and EC+ITC (fractions 22-24). 49-nt and 25-nt transcripts from the first and the second rounds of transcription are indicated by black and red arrows. Note that both complexes have some contamination of PICs (fractions 18-21). **(C)** Eight representative reference-free 2D class averages of EC+EC. **(D)** Same as (C) but for EC+ITC. A large density attributable to TFIIH (indicated by orange arrow heads) is located between EC and ITC. **(E)** A representative 2D class average of EC+EC, with a schematic model, showing the two well-featured densities corresponding to the leading EC and trailing EC, respectively. The density of DNA bridging two polymerases is clearly discernable. **(F)** Same as (E) but for EC+ITC, showing a large density attributable to TFIIH (orange) compared with 2D class averages of EC + EC. The density of DNA is clearly discernable as in (E).

**Figure 5.**
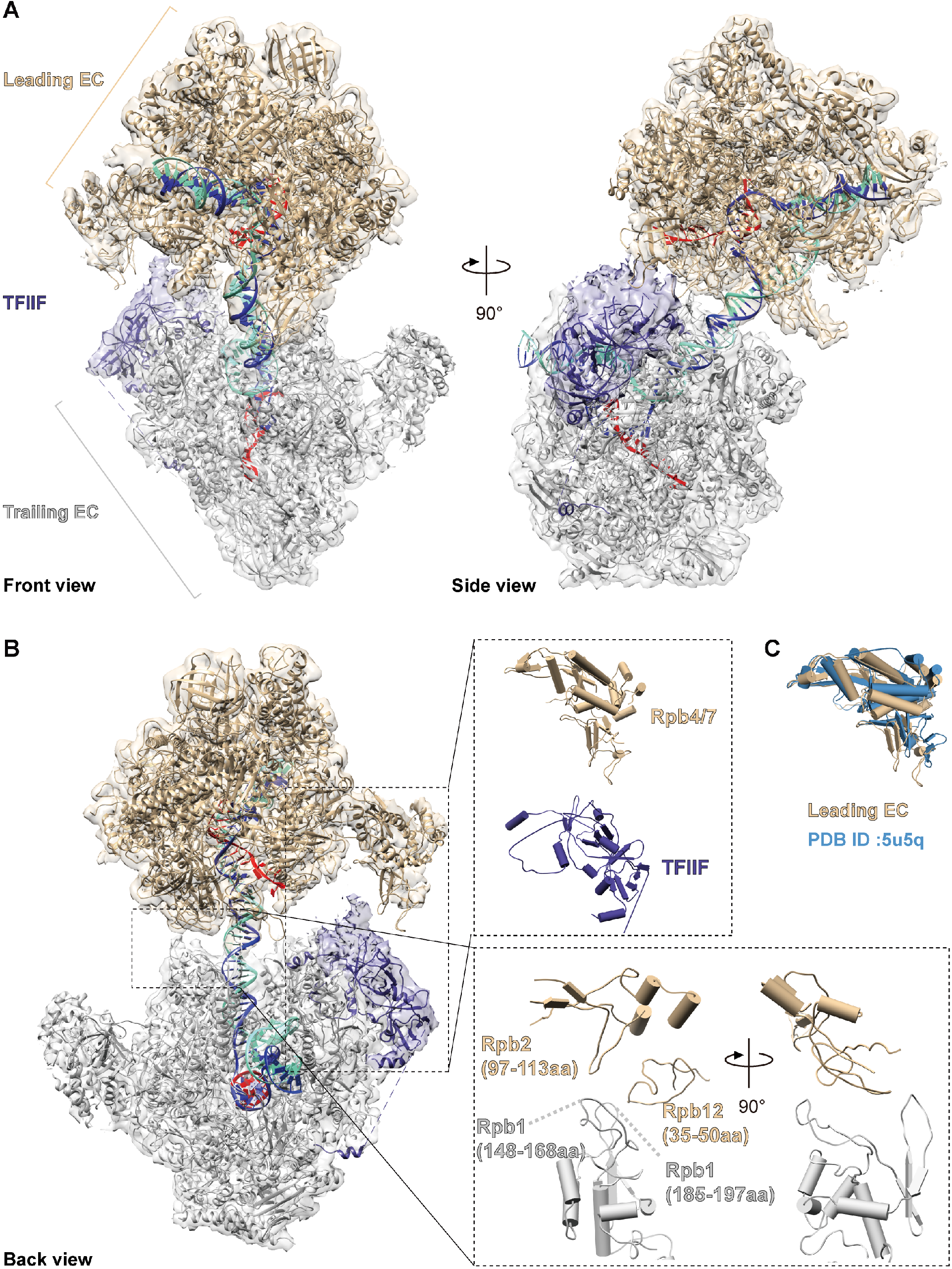
The structure of ECs colliding head-to-end (EC+EC). **(A)** Front (left) and side views (right) of the cryo-EM reconstruction with the model. **(B)** Interactions between two ECs viewed from the back. The Rpb4/7 of the leading EC contacts TFIIF associated with the trailing EC, while Rpb2 and Rpb12 of the leading EC contacts the Rpb1 of the trailing EC. The leading EC, the trailing EC, TFIIF, template DNA, non-template DNA and RNA are colored in tan, gray, navy blue, blue, aquamarine and red, respectively.

In the structure of EC+EC, two colliding ECs span over ~74 bp of DNA (from positions –8 to +66 relative to TSS) (Figure 5). There was a well-ordered density corresponding to TFIIF only on the trailing EC, but not the leading EC (Figure 5A). A previous crystallographic model of an EC complex with a 9-bp DNA-RNA hybrid (PDB ID: 5C4J) (Barnes et al., 2015) was fitted into two corresponding densities with some deviations in the non-template strand of the transcription bubble. Also a previous model of pol II-TFIIF (PDB ID: 5FYW) was fitted without any deviations except a ~10°-rotation of Rpb4/7 subunits of the leading EC, that enabled a direct contact with TFIIF of the trailing EC (Figures 5B-C).

The active site (the nucleotide addition site) of the leading EC was located at the G-stop (+49), while that of the trailing EC was located at +14 (Figures 6A-B, and 6G). This suggests that the trailing pol II that had reached ~+25 to transcribe a ~25-nt RNA, was subjected to extensive (~11bp) backtracking, and arrested at +14. Without this backtracking, two ECs require substantial structural changes in the protein component or/and the DNA component to avoid steric clash at the interface. Consistent with the pol II backtracking, there was density attributed to this backtracked RNA in the funnel of the trailing EC, but not the leading EC (Figures 6C-E). Two-body refinement revealed a ~6°-rotational motion relative to each other, while maintaining the 35-bp spacing between two nucleotide addition sites (Movie S2).

**Figure 6.**
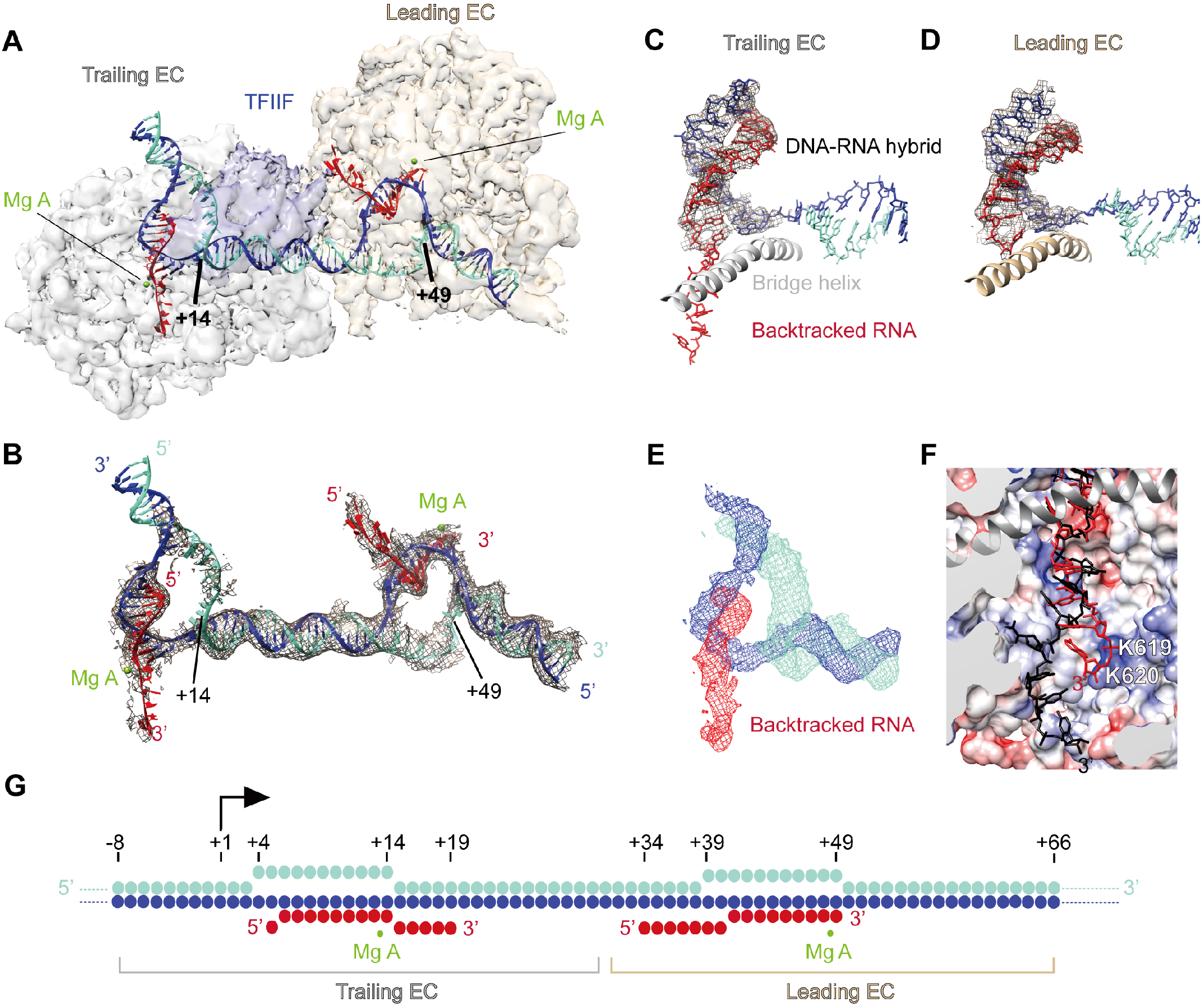
Nucleic acids of ECs colliding head-to-end (EC+EC). **(A)** Unsharpened cryo-EM densities of the trailing EC (gray), the leading EC (tan) and TFIIF (navy blue) are shown as surface, contoured at 2.07 sigma. Template DNA, non-template DNA and RNA are colored in blue, aquamarine and red throughout. Mg A in active center is shown as sphere and colored in green. **(B)** Composite cryo-EM density of nucleic acids. Unsharpened cryo-EM densities are shown as mesh (space gray), contoured at level 2.07 sigma. **(C)** The DNA-RNA hybrid of the trailing EC. Sharpened cryo-EM densities of the DNA-RNA hybrid of the trailing EC are shown as mesh (space gray), contoured at level 4.1 sigma. The bridge helix is shown in gray. **(D)** Same as (C) but for the leading EC, contoured at 3.86 sigma. **(E)** Cryo-EM density of nucleic acids of the trailing EC showing the backtracked RNA (red). Unsharpened cryo-EM densities of nucleic acids are shown as mesh (RNA, red; template DNA, blue; Non-template DNA, aquamarine), contoured at level 1.6 sigma. **(F)** Superposition of the backtracked RNA of the trailing EC (red) with the backtrack site of the arrested Pol II previously determined by crystallography (PDB:3PO2, black). Backtracked RNA in trailing EC shifts towards a positive charged patch that consists of K619 and K620 of Rpb1 in funnel. **(G)** Schematic of nucleic acids. Modelled nucleotides of promoter template are shown with filled circles. TSS (+1) is indicated by black arrow.

Following the substantial backtracking of the trailing EC, specific protein-protein interactions were established at the interface between two ECs (Figure 5B). There were two major points of contact: the first point of contact involved two loops (residues 148-168 and residues 185-197) protruding from Rpb1 clamp of the trailing EC, and the loop of Rpb2 (residues 97-113) and the Rpb12 zinc ribbon (residues 35-50) of the leading EC. The second point of contact involved the tip (residues 134-137) of the dimerization domain of Tfg1 (TFIIF) of the trailing EC, and the tip (residues 125-127) of Rpb7 of the leading EC. Also there were some weak EM densities that were not modeled at the interface. These densities may be attributed to YEATS (residues 1-137) and ET domains (residues 174-244) of Tfg3 as well as the C-terminal WH domain (residues 671-735) of Tfg1 based on an integrative modeling derived from XL-MS (Figure S5).

The template DNA was overall Z-shaped with two kinks at the two active centers of pol II (Figures 6A-B). The 24-bp DNA (from +15 to +38) bridging between two active centers was clearly discerned and modeled with a straight B-form DNA (Figure 6B). The density of the DNA-RNA hybrid in each active center was traceable (Figures 6B-D): in the leading EC, 16 ribonucleotides of the 49-nt transcript were visualized: 9 ribonucleotides from the 3’ end formed a hybrid with the template DNA (positions from +41 to +49), with the 3’ end of the transcript (3’-O-methyl guanosine 5′-monophosphate) being aligned at the nucleotide addition site (designated i+1 position) in the active center, while a stretch of adjacent seven ribonucleotides was in the RNA exit tunnel (Figures 6A-B and 6G). In the trailing pol II, 15 ribonucleotides of the ~25-nt transcript were discernible (Figures 6C and 6G). 9 ribonucleotides formed a hybrid with the template DNA (positions from +6 to +14) in the active site, and adjacent five ribonucleotides of the backtracked RNA were observed in the pol II pore and funnel (Figure 6E): two ribonucleotides of the backtracked RNA at positions i+2 and i+3 were in the pore as previously observed by X-ray crystallography(Wang et al., 2009), while three ribonucleotides at positions i+4, i+5, and i+6 lie on a positively charged patch composed of Lys619 and Lys620 of Rpb1 in the funnel, not observed in any previous transcribing complex structures (Figure 6F). The path of the backtracked RNA differed from that of the backtrack site previously observed by X-ray crystallography (Figure 6F) (Cheung and Cramer, 2011). The backtracked RNA is nonetheless incompatible with TFIIS, and must be displaced from these sites for TFIIS-induced transcription resumption from the backtracked state (Cheung and Cramer, 2011).

Two lines of evidence support the specificity and functional significance of the EC+EC complex. First, previous exonuclease footprinting of elongation complexes without TFIIF, exhibited greater variability in the distance between two ECs upon head-to-end collision as well as much more extensive backtracking of the trailing EC (~50 bp backtracking upon encountering a leading EC)(Saeki and Svejstrup, 2009), supporting the specificity of the EC+EC conferred by TFIIF. Second, the trailing EC stalled at +14 completed promoter escape, whereas initiation complexes stalled at any positions before +27 failed to escape promoter in our single round transcription system (Fujiwara et al., 2019). The leading EC stalled at +49 from a preceding round of transcription likely plays a positive role in promoter escape of a trailing ITC, rather than simply acting as a roadblock of TFIIH translocation.

### Promoter escape of the ITC is facilitated by a transcribing pol II at promoter proximal regions

Direct support for the role of the leading EC in promoter escape of the trailing ITC came from cryo-EM analysis of the EC+ITC (Figures 4D and 4F). All 2D class averages showed a large (~500 kDa) density attributable to TFIIH in a space between two polymerases (indicated by orange arrow heads in Figure 4D). The assignment of TFIIH was further validated by a comparison with a 2D projection from a 3D model of EC+core ITC (ITC lacking TFIIH) (Figure S7B). Of the eight class averages we obtained, the top four populated classes maintained a similar spacing between the EC and the ITC as in the EC+EC, through the direct protein-protein interactions described above (upper row in Figure 4D). In these class averages, the DNA double helix was accommodated in the pol II downstream cleft of the trailing ITC, while TFIIH was dissociated from the DNA and displaced from its position in the ITC of the G-less26 complex (schematically illustrated in Figure 4F, see also Figure S7A). This conformational change of the ITC, as an irreversible critical transition from initiation to elongation (see Discussion), was evidently facilitated by the presence of the leading EC. In the other classes, two ECs were apparently separated from each other, suggesting that the trailing ITC was subjected to more extensive backtracking (lower row in Figure 4D), which may further require TFIIS for transcription resumption from the backtracked state.

## Discussion

Structural and mechanistic studies of transcription initiation involving TFIIH have been hampered by poor efficiency of initiation reaction in vitro (commonly ~0.01-0.1 transcripts per PIC). Previous structural models of transition from initiation to elongation were derived from complexes with artificially open templates, and not obtained by the catalytic activity of TFIIH. Thus how TFIH directs promoter melting, TSS scanning, and promoter escape (Dvir et al., 1997; Fishburn et al., 2016; Luse, 2013; Qiu et al., 2020; Spangler et al., 2001) remains to be resolved.

To dispel this long-standing mystery of the transcription initiation process, we have developed a highly efficient in vitro reconstitution from the yeast at quality and quantity amenable to structure determination (Fujiwara and Murakami, 2019; Murakami et al., 2013a). As a notable achievement reported here, we have arrived at a complete description of pol II transcription from initiation by the 33-subunit PIC through promoter escape, to finally reach elongation. Our structural data provide a direct evidence that the ITC retaining all GTFs continues until it encounters an EC at promoter proximal regions from a preceding round of transcription, and that two polymerases occlude TFIIH binding to facilitate promoter escape. Promoter escape, viewed in the past as no more than dissociation of pol II from promoter, now appears mechanistically varied, with important regulatory consequences.

Three distinct forms of the PIC were defined in this study: relative to PIC1 in a form similar to previous structures (Dienemann et al., 2019; Murakami et al., 2015b), PIC2 and PIC3 exhibited ~20 Å and ~30 Å shifts in the location of TFIIH, and repositioning of Ssl2 on DNA by one turn of dsDNA, along with greater degrees of DNA distortion. In PIC1, the location of TFIIH is constrained primarily by the contact with the relatively straight and rigid double-stranded DNA. Upon the DNA distortion in PIC2/PIC3, the positional constraint of TFIIH imposed by the double-stranded DNA is relieved, such that TFIIH is rather stabilized by direct protein-protein contacts with TFIIE. Locations of GTFs of the PIC3 closely correspond to those in the ITC, indicating the functional significance of PIC3, as well as PIC2, as intermediates on a path to the open complex formation. However, apparently a transition from PIC1 to PIC2/PIC3 requires rebinding of TFIIH on DNA, due to the upstream shift in the location of Ssl2 on DNA. Energy barriers required for this rebinding may indicate some functional differences between PIC2/PIC3 and PIC1.

The possible functional differences between distinct forms of the PIC may relate to two forms of TFIIH: only the weak-binding state, which most likely represent a post-translocation state of TFIIH, was exclusively identified in PIC2/PIC3, whereas the strong-binding state (pre-translocation state) was apparently favored in PIC1. This suggests that locations of TFIIH in PICs would shift the conformational equilibrium among coexisting translocation states to regulate translocase activity of TFIIH. Although previous and current structures of TFIIH did not directly resolve nucleotide states, there is a consensus observation that the strong-binding state (pre-translocation state) was exclusively identified from specimens with a non-hydrolysable ATP analogue or without ATP, while the weak-binding state (post-translocation state) was identified only when ATP was provided (this study and (van Eeuwen et al., 2021b)).

In previous structures of the open PIC (He et al., 2016; Schilbach et al., 2017), downstream dsDNA was stably accommodated in the pol II downstream cleft and the further downstream end was simultaneously bound by TFIIH. Considering that these models resemble those from other transcription systems devoid of equivalent translocases and that a similar open complex could be formed in the absence of TFIIH (Plaschka et al., 2016), they may represent the pathway to TFIIH-independent transcription (Alekseev et al., 2017; Holstege et al., 1995; Parvin and Sharp, 1993). By contrast, in our bona fide ITC, TFIIH precluded such stable accommodation of the downstream DNA in the pol II downstream cleft, and directed open complex formation that were essentially maintained by GTFs, but not pol II. Due to the lack of the direct contact with the downstream DNA, pol II may have a degree of freedom of lateral movement along the template, and thus confer TFIIH-dependent properties in TSS utilization, initial RNA synthesis, and promoter escape (Bradsher et al., 2000; Dvir et al., 1997; Fishburn et al., 2016; Fujiwara et al., 2019; Murakami et al., 2015a; Spangler et al., 2001).

Previous biochemical and biophysical data suggest that a bona fide ITC is long-persisting and that promoter escape occurs after synthesis of dozens of nucleotides (Fazal et al., 2015; Fujiwara et al., 2019; Luse, 2019). However this is unlikely to occur in cells as the initially transcribing pol II is thought to encounter another pol II or a nucleosome at promoter proximal regions shortly after the initiation of transcription (Ehrensberger et al., 2013). Thus our G-less 49 complex may represent a more complete picture of promoter escape occurring in vivo, as an ITC is formed in the presence of EC stalled at +49 from a preceding round of transcription (Figure 7). Contrary to expectation, an EC at promoter proximal regions supported a positive role in transcription rather than acting as a transcriptional roadblock; TFIIH of the ITC was occluded between two transcribing polymerases, followed by partial dissociation of TFIIH (resulting in EC+ITC) or complete dissociation of TFIIH (resulting in EC+EC) (stage 3 or 4 in Figure 7). In the EC+EC, the trailing pol II completed promoter escape after transcribing ~25 nt RNA, while in the EC+ITC, the trailing ITC apparently failed to escape promoter, but successfully accommodated the downstream DNA in the pol II downstream cleft (upper row of Figure 4D). Both structures markedly contrast to the G-less 26 ITC that failed to escape promoter in the absence of such EC in front of it (Figure 3). Although previous biochemical data suggest that the 8-9-bp DNA-RNA hybrid is the minimum requirement for the formation of pol II-DNA-RNA complex (Kireeva et al., 2000), we posit the interaction between pol II and the downstream DNA confers additional stability to keep the polymerase in register at the 3’-end of RNA. Before this transition, TFIIH continuously draws dozens of DNA base pairs by scrunching (Fazal et al., 2015; Tomko et al., 2017), on which initially transcribing pol II (as well as TSS scanning pol II) has a degree of freedom of lateral movement along the template. Therefore the stable accommodation of the downstream DNA in the pol II downstream cleft, which marks the end of the requirement for TFIIH, represents an irreversible critical transition from initiation to elongation. This explains why the G-less 27 complex stalled at +27 released transcripts by extensive backtracking of pol II upon removal of ATP during gradient sedimentation, whereas the G-less 49 complex completely retained transcripts of similar length (~25 nt or shorter) in the trailing ITC (Fujiwara et al., 2019). It should be noted that even after the entry of downstream DNA into the pol II cleft, in some ITCs, GTFs remained bound to pol II (Figure 4D), which may further require additional elongation factors such as the capping enzyme and/or Spt4/5 to displace TFIIE from pol II (Fujiwara et al., 2019).

**Figure 7.**
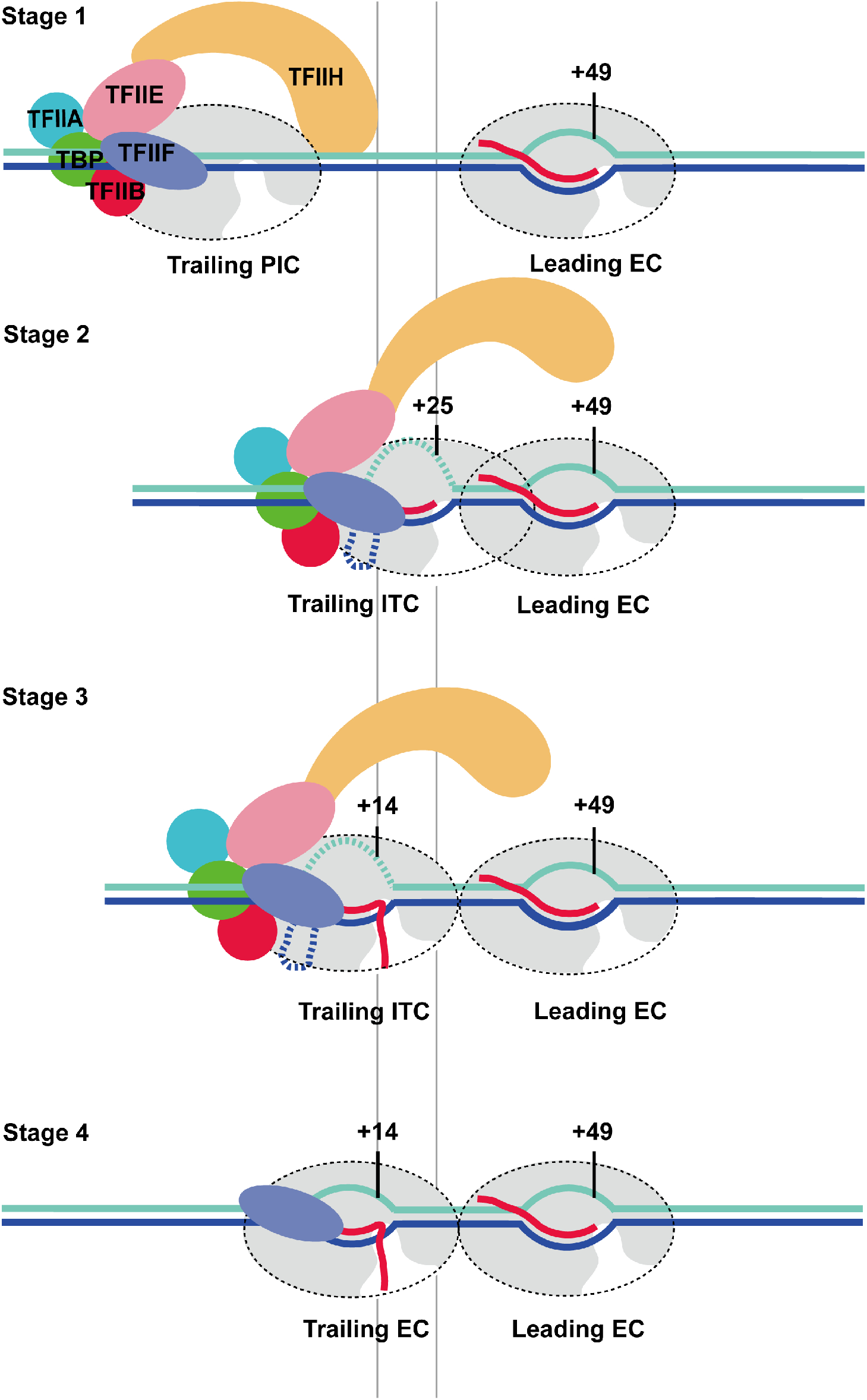
Schematic of promoter escape facilitated by the leading EC. A PIC is assembled on promoter, while an EC is stalled at promoter proximal regions on the G-less 49 template (Stage 1). PIC reels ~80 bp of downstream DNA by scrunching and initiates transcription, while retaining fixed upstream contact within the complex. Initially-transcribing pol II in the trailing ITC encounters the leading EC stalled at +49 and occludes TFIIH binding (stage 2). The trailing EC is backtracked by ~11bp and arrested at +14 (EC+ITC, Stage 3), followed by dissociation of GTFs and bubble collapse (EC+EC, Stage 4).

Lastly, when the structures of pol II (EC)-DSIF-NELF complex (Vos et al., 2018), in a canonical form of prompter-proximal paused pol II in mammalian systems, is aligned with the leading EC of the EC+EC complex, there is no steric clash of the trailing EC with NELF, but partial clash with DSIF. Also Mediator, which serves a critical role in promoter escape (Jeronimo and Robert, 2014; Takahashi et al., 2011; Wong et al., 2014), has no steric clash with the leading EC when the trailing EC is aligned with the PIC-Mediator complex (Robinson et al., 2016). It will be of great interest to pursue possible positive/negative regulations of promoter escape by such general factors and determine the underling structural basis.

## Acknowledgments

We thank Dr. Sudheer Molugu and Electron Microscopy Resource Laboratory at the University of Pennsylvania for use of equipment and assistance with cryo-EM sample screening. We thank the University of Massachusetts CryoEM Core Facility, Drs. Chen Xu, KangKang Song and Kyounghwan Lee for their constant support and assistance in data collection of G-less 26 complexes. We thank Thomas Edwards, Adam Wier at the National Cancer Institute’s National Cryo-EM Facility at the Frederick National Laboratory for Cancer Research for data collection of the G-less 49 complex (EC+EC). We thank Janette Myers, Nancy Meyer, and Harry Scott at the Pacific Northwest Center for Cryo-EM (PNCC) for cryo-EM screening and data collection of G-less 49 complexes (EC+ITC). We thank members in the lab for helpful discussions and suggestions.

## Author contributions

C.Y. and R.F. conducted the experiments, C.Y., R.F., S.S., and K.M. performed cryo-EM image analysis, C.Y. and H.J.K. performed cross-linking mass spectrometry (XL-MS), H.J.K. and B.A.G. analyzed XL-MS data, J.J.G performed integrative modeling. C.Y., R.F., and K.M. prepared figures, designed the study, and wrote the manuscript.

## Data availability

Cryo-EM and model data: The electron density reconstructions ad final models are deposited with the Electron Microscope Data Base (EMDB) under accession codes EMD-23904 (PIC1), EMD-23905 (PIC2), EMD-23906 (PIC3), EMD-23907 (TFIIH weak binding state), EMD-23908 (ITC), and EMD-23789 (EC+EC), and with the Protein Data Bank (PDB) under accession codes 7ML0 (PIC1), 7ML1 (PIC2), 7ML2 (PIC3), 7ML3 (TFIIH weak binding state), 7ML4 (ITC), and 7MEI (EC+EC).

XL-MS data: Crosslinking Mass-Spectrometry data of EC+EC was deposited in PXD025758.

